# Calcium/calmodulin-dependent protein kinase II regulates mammalian axon growth by affecting F-actin length in growth cone

**DOI:** 10.1101/518076

**Authors:** Feng Xi, Ren-Jie Xu, Jin-Hui Xu, Wei-Hua Wang, Jin-Jin Ma, Feng Wang, Yan-Xia Ma, Shi-Bin Qi, Jing-Cheng Zhang, Hao-Nan Zhang, Xu-Zhen Qin, Jian-Quan Chen, Bin Li, Chang-Mei Liu, Hui-Lin Yang, Bin Meng, Saijilafu

**Affiliations:** Department of Orthopaedics, the First Affiliated Hospital, Orthopaedic Institute, Soochow University, Suzhou, P.R.China, 215007; Department of Orthopaedics, Suzhou Municipal Hospital/the Affiliated Hospital of Nanjing Medical University, Suzhou, Jiangsu, China; State Key Laboratory of Stem Cell and Reproductive Biology, Institute of Zoology, Chinese Academy of Science, Beijing, China; Savaid Medical School, University of Chinese Academy of Sciences, Beijing, China

**Keywords:** axon growth, CaMKII, sensory neurons, cytoskeleton, F-actin

## Abstract

While axon regeneration is a key determinant of functional recovery of the nervous system after injury, it is often poor in the mature nervous system. Influx of extracellular calcium (Ca^2+^) is one of the first phenomena that occur following axonal injury, and calcium/calmodulin-dependent protein kinase (CaMKII), a target protein for calcium ions, regulates the status of cytoskeletal proteins such as F-actin. Herein, we found that peripheral axotomy activates CaMKII in dorsal root ganglion (DRG) sensory neurons, and inhibition of CaMKII impairs axon growth in both the peripheral and central nervous systems (PNS and CNS, respectively). Most importantly, we also found that activation of CaMKII promotes PNS and CNS axon growth, and regulatory effects of CaMKII on axon growth occur via affecting the length of the F-actin. Thus, we believe our findings provide clear evidence that CaMKII is a critical modulator of mammalian axon regeneration.

## Introduction

Failure of axonal regeneration in the damaged mammalian nervous system is the main impediment to its functional recovery. However, injured axons of the adult central nervous system (CNS) cannot regenerate spontaneously owing to their poor capacity of intrinsic axon growth and the inhibitory microenvironment. Compared to the CNS neurons, axons of the peripheral nervous system (PNS) neurons can partly regenerate after injury; however, its functional outcome is often poor, especially in humans, resulting in proximal lesions. This is largely because of atrophic changes in chronically denervated Schwann cells in distal nerves and target atrophy [1]. Therefore, the rate of axon growth is a major limitation in functional recovery of the injured nervous system. Understandably, most studies concerning axon regeneration in the past few decades have focused on ways to antagonize the effects of inhibitory molecules and/or activate the intrinsic axon growth capacity of adult neurons [2–6]. However, thus far, little attention has been paid to enhance the rate of axon regeneration by modulating the local axonal cytoskeleton assembly at the growth cone—where axon growth actually occurs. Additionally, increasing evidence has shown that long-distance axon regeneration is achieved by regulating the cytoskeletal elements at the nerve growth cone [7–11]. Thus, modulation of the cytoskeletal protein assembly in the growth cone may be an effective strategy to promote axon regeneration in the damaged nervous system.

Calcium/calmodulin-dependent protein kinase II (CaMKII) is the most conventional member of the CaMK family that regulates a series of functions in the nervous system, such as neurotransmitter synthesis/release, synaptic plasticity, learning, and memory. Four isoforms of CaMKII are expressed in the nervous system: α, β, γ, and δ; among these, the γ isoform is most abundantly expressed in the dorsal root ganglion (DRG) neurons; while isoforms α and β are primarily expressed in the brain [12]. Calcium/calmodulin can bind to CaMKII and activate it through autonomous phosphorylation and subsequently sustain its kinase activity [13]. A previous study demonstrated that calcium influx occurred immediately after axonal damage; furthermore, this calcium influx plays key roles subsequent to axonal growth [14]. In addition, another study showed that overexpression of CaMKII promotes neurite outgrowth from neuro2a and NG108-15 neuroblastoma cells [15]. Moreover, CaMKII activity is also specifically required for NCAM, and N-cadherin-induced neurite outgrowth [16]. Thus, it is likely that CaMKII protein could regulate mammalian axon regeneration; however, to our best knowledge, the functional role of CaMKII on primary neuronal axon regeneration has not yet been studied.

Herein, we found that the phosphorylation level of CaMKII in DRG sensory neurons was increased because of peripheral nerve injury. Moreover, the pharmacological inhibition of CaMKII activity or genetic knockdown of CaMKIIγ with a specific siRNA blocks axon growth in DRG sensory neurons and developing CNS neurons. By contrast, the pharmacological activation of CaMKII with CdCl_2_ enhances the axon growth of DRG sensory neurons and developing CNS neurons. Additionally, we found that inhibition of CaMKII activity reduces the F-actin length in the growth cone, and the activation of CaMKII increases the F-actin length. Taken together, these findings provide clear evidence that CaMKII is a critical modulator of mammalian axon regeneration.

## Results

### 1. Peripheral axotomy increases the phosphorylation level of CaMKII in DRG neurons

Previous studies have shown that axonal injury induces Ca^2+^ influx at the lesion site [17], and calcium/calmodulin binds to CaMKII to activate it by autonomous phosphorylation. Thus, we first examined the phosphorylation levels of CaMKII in DRG neurons after sciatic nerve axotomy. Immunohistochemical staining performed three days after the nerve injury showed elevated phosphorylation levels of CaMKII in the DRG neurons compared to the naïve DRG neurons (Fig. 1A). In addition, western blot data also further confirmed that the level of phosphorylated CaMKII protein was also increased after sciatic nerve injury (Fig. 1B, C), indicating that peripheral axotomy activates the CaMKII signaling pathway in peripheral sensory neurons.

**Figure 1:**
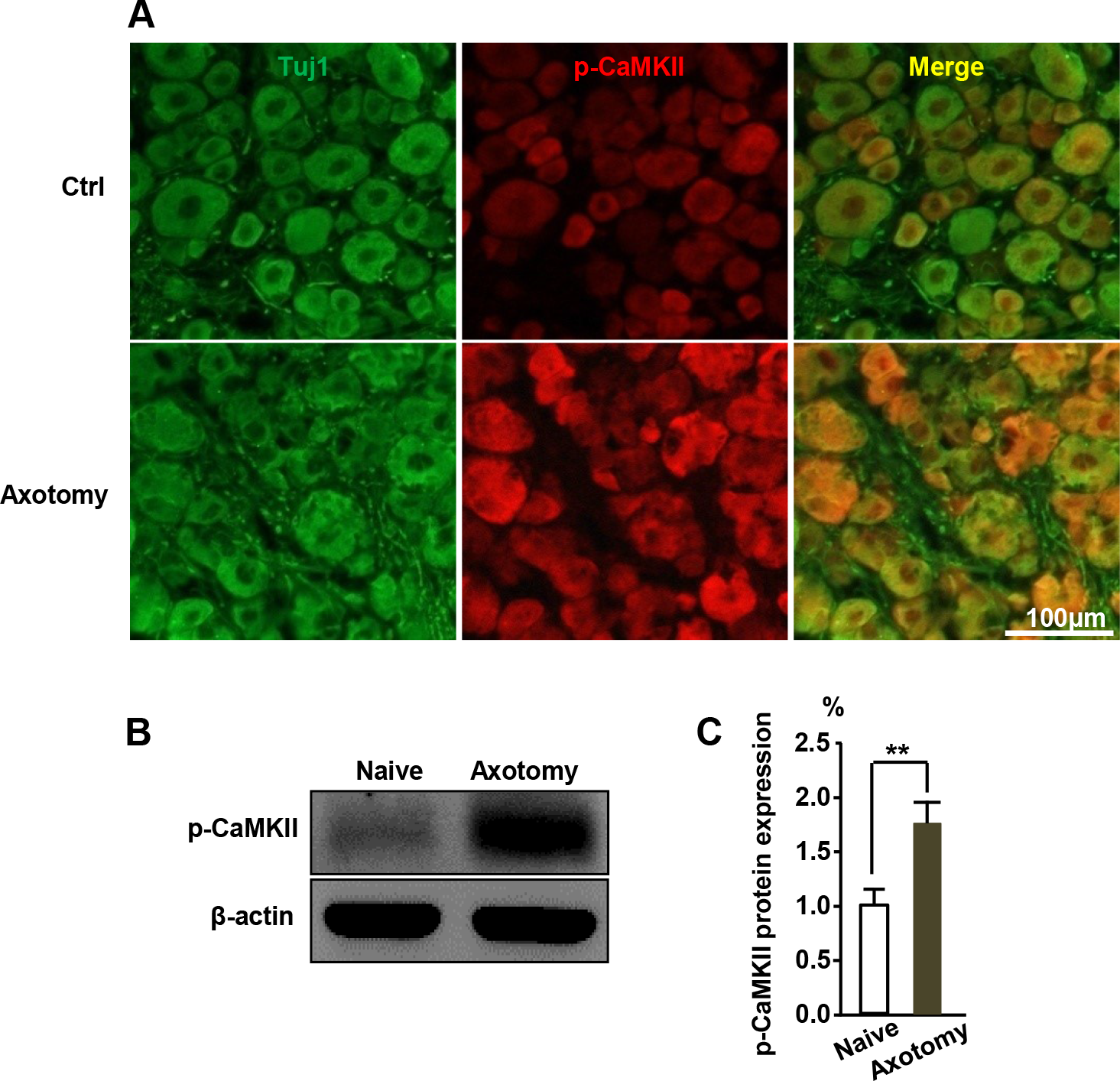
Peripheral axotomy increases phospho-CaMKII expression in DRG neurons. (A) Compared to the naïve control group, p-CaMKII expression was increased in the DRG neurons after sciatic nerve axotomy. Green represents TUJ1 and red, p-CaMKII. Scale bar: 100 μm. (B) Results of western blotting showing increased p-CaMKII expression levels in DRGs seven days after sciatic nerve axotomy. (C) Quantification of p-CaMKII western blot band (normalized to actin, n = 3).

### 2. Inhibition of CaMKII activity prevents axon growth from adult DRG neurons

Next, we investigated the functional role of CaMKII on axon regeneration. Dissociated DRG neurons from adult mice were treated with DMSO and either 2.5 or 5.0 μM KN93—a specific CaMKII inhibitor—and cultured for 3 days. When the phosphorylation level of CaMKII was effectively blocked by KN93 (Fig. 2A, B), the regenerative axon growth from KN93 treatment group was significantly shorter than the control group (Fig. 2C, D). To confirm the pharmacological results, we knocked down CaMKIIγ expression using specific siRNAs in the cultured DRG sensory neurons. Among four isoforms of CaMKII, the γ isoform is abundantly expressed in DRG neurons. Our western blot results also showed that siRNA against the γ isoform efficiently suppressed CaMKII expression in the DRG neurons (Fig. 2E, F). Furthermore, consistent with pharmacological results, gene silencing of CaMKIIγ expression with specific siRNA dramatically inhibited regenerative axon growth of DRG neurons (Fig. 2G, H).

**Figure 2:**
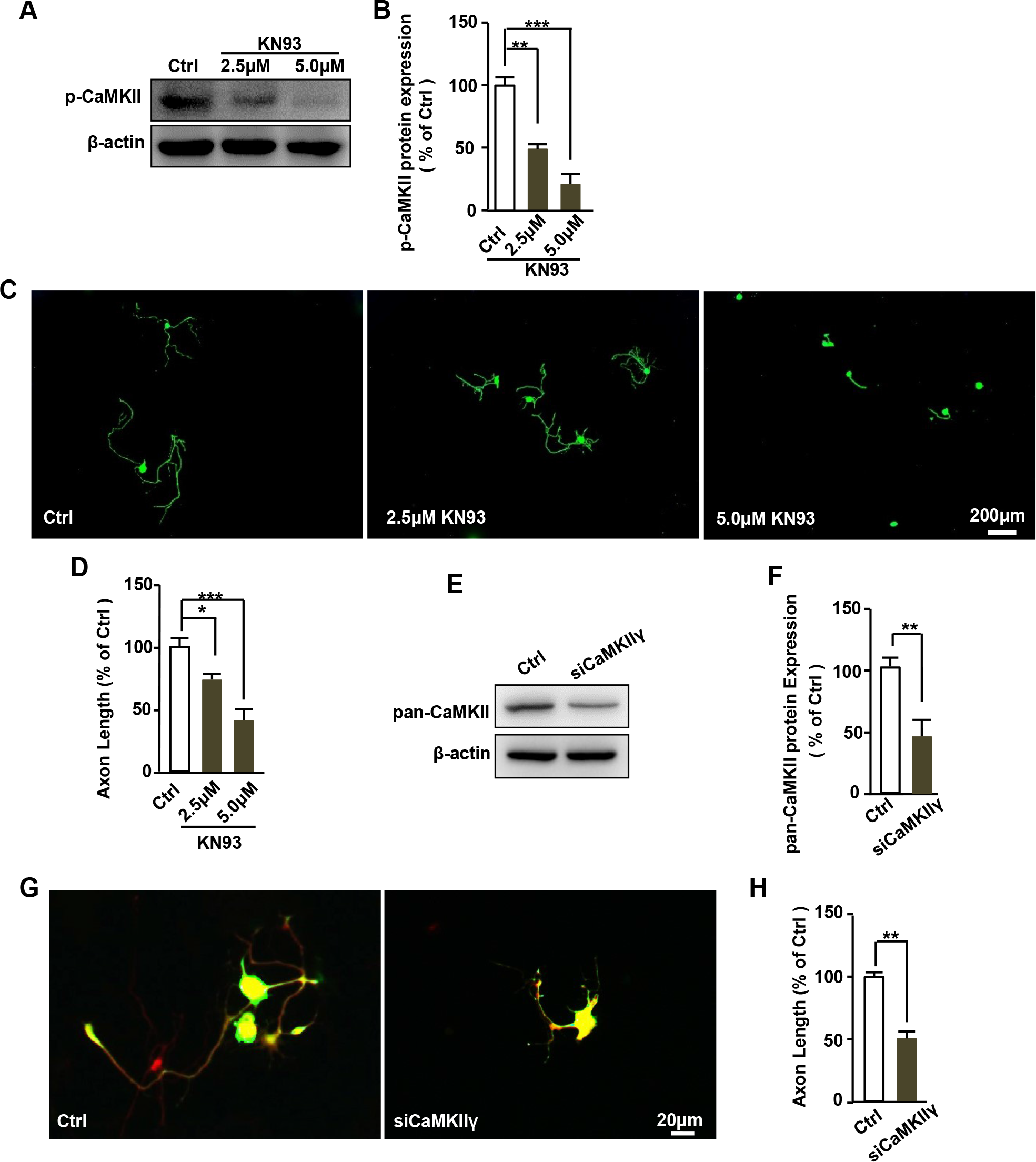
Inhibition of CaMKII activity prevents axon growth in adult DRG neurons. (A) Inhibition of CaMKII activity with its pharmacological inhibitor KN93 significantly blocks phosphorylation level of CaMKII in DRG neurons. (B) Quantification of western blotting of (A). (C) KN93 treatment markedly inhibits the DRG neuronal axon growth in vitro in a dose-dependent manner. Neurons were stained with TUJ1 antibody, Scale bar: 200 μm. (D) Quantification of the average axon length from three independent experiments (n=3). (E) A specific siRNA against CaMKIIγ significantly blocks pan-CaMKII expression in the DRG neurons. (F) Quantification of siCaMKIIγ knocking down efficiency in (E). (G) Transfection of siCaMKIIγ markedly inhibits DRG neuronal axon growth. Scale bar: 20 μm. (H) Quantification of the average axon length from three independent experiments (n=3).

### 3. CaMKIIγ knockdown inhibits peripheral sensory axon regeneration in vivo

To further explore the roles of CaMKII on axon regeneration in vivo, we used in vivo DRG electroporation technique to introduce si-CaMKIIγ plus EGFP into adult mouse lumbar (L) 4/5 DRG neurons. Two days later, the ipsilateral sciatic nerve was crushed using fine forceps. As described in our previous study, three days after the sciatic nerve injury, the axon regeneration was assessed in whole mount flattened nerve [18]. All EGFP-labeled axons were then manually traced from the lesion site to the distal axonal tip to measure the accurate length of the regenerating axon. Consistent with our in vitro results, the axon regeneration was significantly impaired by siCaMKIIγ (Fig. 3A-C), thereby indicating that the activity of CaMKII is required for peripheral axon regeneration in vivo.

**Figure 3:**
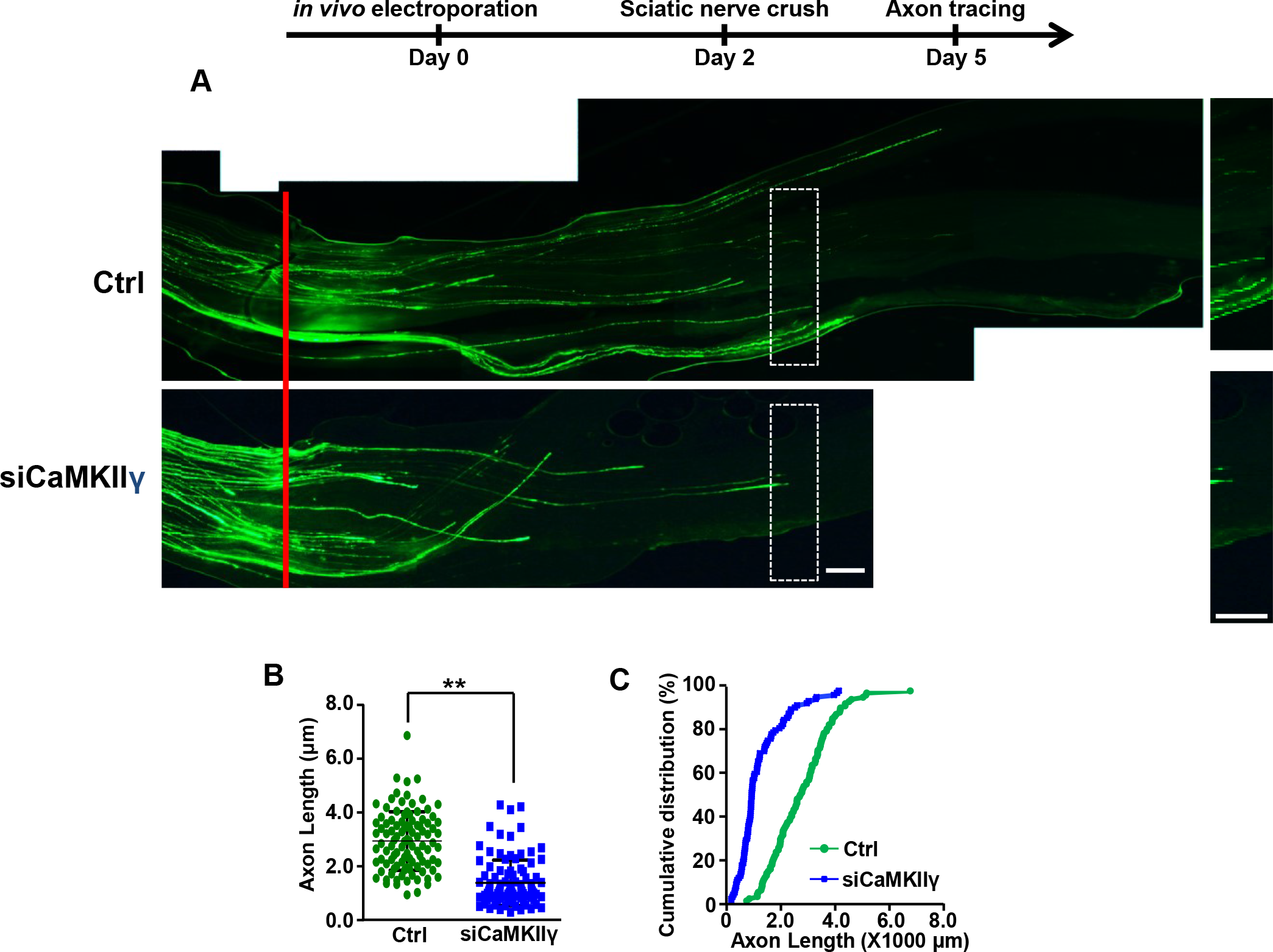
CaMKII activity is required for the peripheral sensory axon regeneration in vivo. (A) Representative images of axon regeneration in vivo following siCaMKIIγ transfection. Red arrowheads indicate the crush sites. Scale bar: 200 μm. (B, C) Quantification of average length of EGFP-positive regenerating axons showing CaMKIIγ knock-down in DRG neurons significantly inhibits axon regeneration in vivo (n = 6 mice for each).

### 4. CaMKII activity regulates axon growth of embryonic cortical and hippocampal neurons

Typically, intrinsic axon regeneration ability of the CNS and PNS neurons is different. Therefore, we questioned whether CaMKII regulates the axon regeneration ability of CNS neurons. To test this hypothesis, we treated E14.5 cortical and E18 hippocampal neurons with DMSO or 2.5 μM KN93 for three days. Similarly, treatment of KN93 induced significant axon growth inhibition in both cortical and hippocampal neurons (Fig. 4A-D). To further confirm the pharmacological inhibitor results, we used a specific siRNA against CaMKIIγ to knock down CaMKIIγ expression in the E14.5 cortical and E18 hippocampal neurons. In line with the results of pharmacological inhibition results, siCaMKIIγ transfection also markedly inhibited both cortical and hippocampal neuronal axon growth (Fig. 4E-H). Taken together, these results suggest that CaMKII activity is required for CNS axon growth as well.

**Figure 4:**
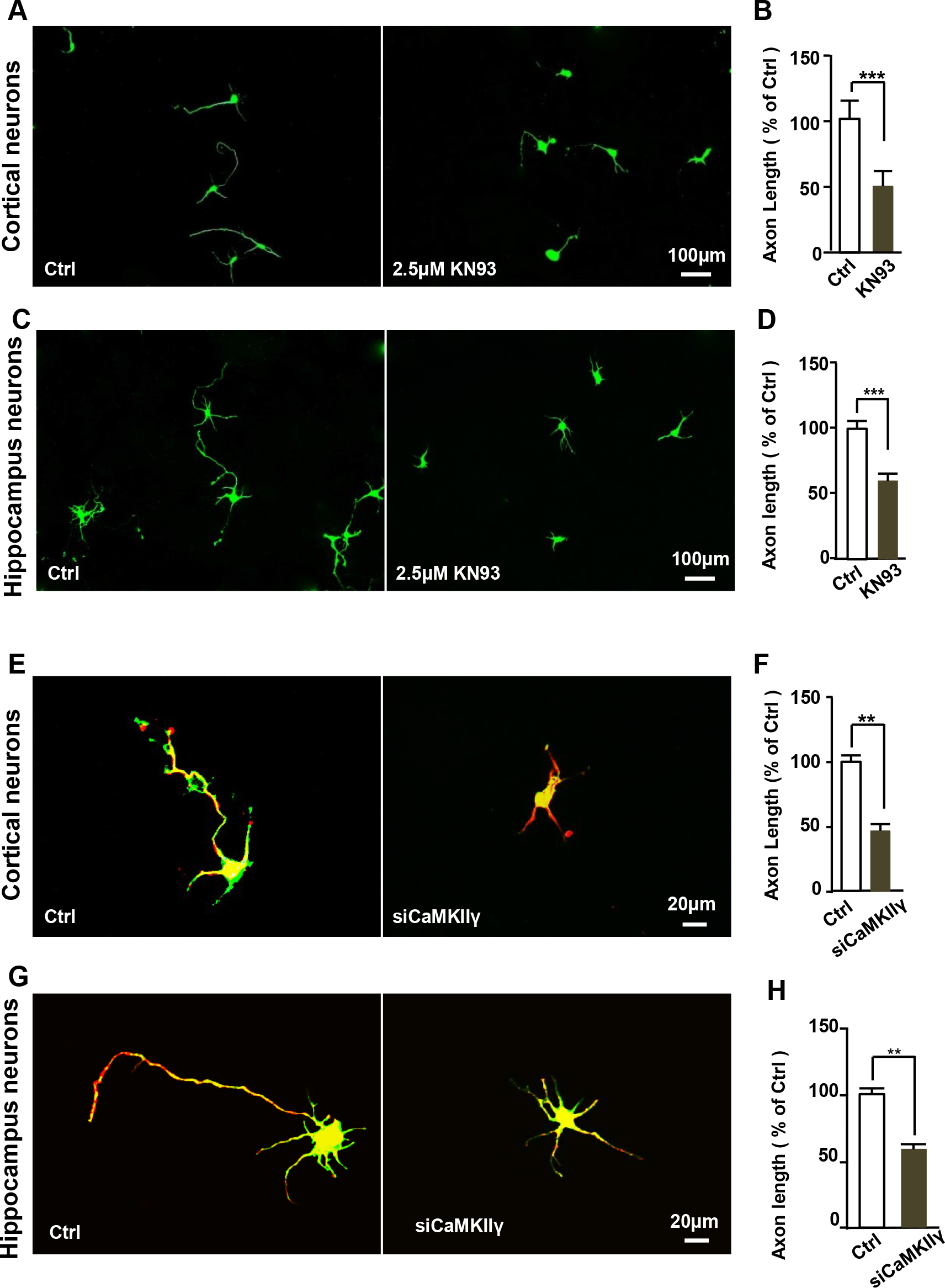
Inhibition of CaMKII activity impairs axon growth in embryonic CNS neurons. (A) Representative images showing that inhibition of CaMKII activity with the pharmacological inhibitor—2.5 μM KN93—blocked cortical neuron axon growth. Scale bar: 100 μm. (B) Quantification of the average axonal lengths of cortical neurons shown in (A) from three independent experiments. (C) Representative images showing inhibition of CaMKII activity with the pharmacological inhibitor—2.5 μM KN93—blocks hippocampal neuronal axon growth. Scale bar: 100 μm. (D) Quantification of the average axonal lengths of hippocampal neurons shown in (C) from three independent experiments. (E) Representative images showing knockdown of CaMKII activity with a specific siCaMKIIγ blocks cortical neuron axon growth. Scale bar: 20 μm. (F) Quantification of the average axonal lengths of cortical neurons shown in (E) from three independent experiments. (G) Representative images showing knockdown of CaMKII activity with a specific siCaMKIIγ blocks hippocampal neuron axon growth. Scale bar: 20 μm. (H) Quantification of the average axonal lengths of hippocampus neurons shown in (G) from three independent experiments.

### 5. Activation of CaMKII with CdCl2 promotes axon growth in adult DRG neurons and embryonic CNS neurons

A previous study showed that cadmium ions could activate CaMKII in mesangial cells [19]. Consistent with that, we too found that treatment with 10.0 μM CdCl_2_ significantly increased the expression level of phosphorylated CaMKII in cultured DRG neurons (Fig. 5A, B). Hence, we further examined whether activation of CaMKII with CdCl_2_ regulates the vitro axon growth. Interestingly, our results showed that CdCl_2_ could significantly promote the axon growth in adult DRG neurons. Similarly, we also found that treatment of CdCl_2_ promotes embryonic cortical (Fig. 5E, F) and hippocampal neuronal axon growth (Fig. 5G, H). Thus, these results indicate that activation of CaMKII sufficiently promotes neuronal axon growth in the adult PNS and developing CNS, in vitro.

**Figure 5:**
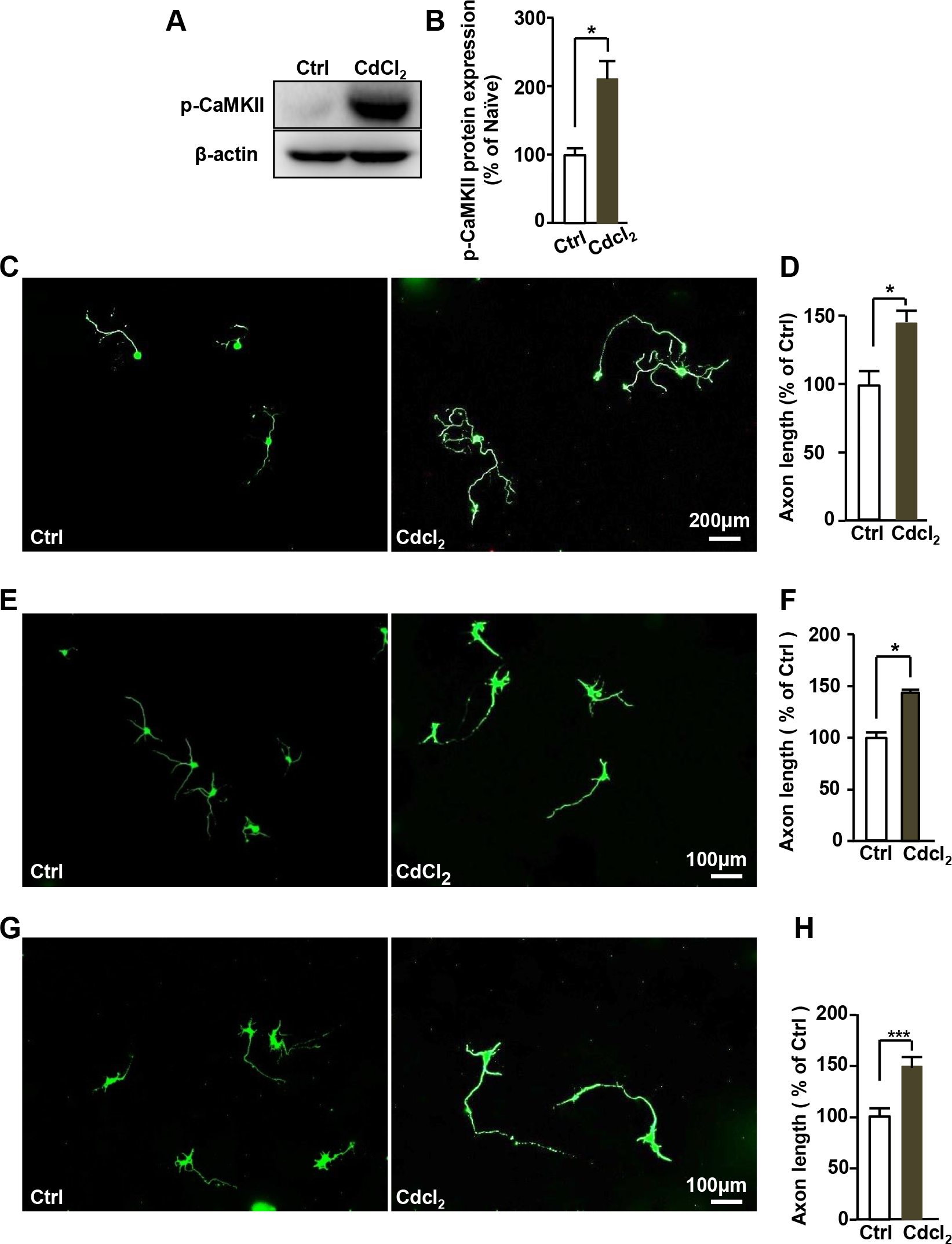
Activation of CaMKII activity promotes axon growth in adult DRG and developing CNS neurons. (A) Western blot images showing that phospho-CaMKII expression level is increased in adult DRG neurons after administration of 10.0 μM CdCl_2_. (B) Quantification of p-CaMKII level in (A). (C) Activation of CaMKII with 10.0 μM CdCl_2_ promotes adult DRG neuronal axon growth in vitro. Scale bar: 200 μm. (D) Quantification of the average axonal lengths of adult DRG neurons shown in (C) from three independent experiments. (E) Activation of CaMKII with 10.0 μM CdCl_2_ promotes axon growth of embryonic cortical neurons. All neurons were stained with TUJ1 antibody. Scale bar: 200 μm. (F) Quantification of the average axonal lengths of adult DRG neurons shown in (E) from three independent experiments. (G) Activation of CaMKII with 10 μM CdCl_2_ promotes axon growth of embryonic hippocampal neurons. Scale bar: 200 μm. (H) Quantification of the average axonal lengths of adult DRG neurons shown in (G) from three independent experiments.

### 6. Activation of CaMKII with CdCl2 prevents injury-induced axon retraction of the cortical spinal tract

Axons of the cortical spinal tract retract in response to spinal cord injury, and subsequently form retraction bulbs [20–23]. Our cell culture results indicate that activation of CaMKII with CdCl2 promotes axon growth in embryonic CNS and adult PNS. Thus, we further examined whether activation of CaMKII with CdCl_2_ promotes in vivo axon regeneration by using a spinal cord injury model. After spinal cord crush injury was established at the T8 level, 1.0 mg/kg CdCl_2_ was intraperitoneally injected every day for 4 weeks. We found that injection of CdCl_2_ could prevent injury-induced axon retraction of the cortical spinal tract (Fig. 6A, B). Unfortunately, however, we did not observe any regenerating axons beyond the lesion site in the spinal cord. Thus, although these results indicate that activation of CaMKII could prevent injury-induced axon retraction, it is difficult to further promote axon regeneration across the lesion site in the injured spinal cord.

**Figure 6:**
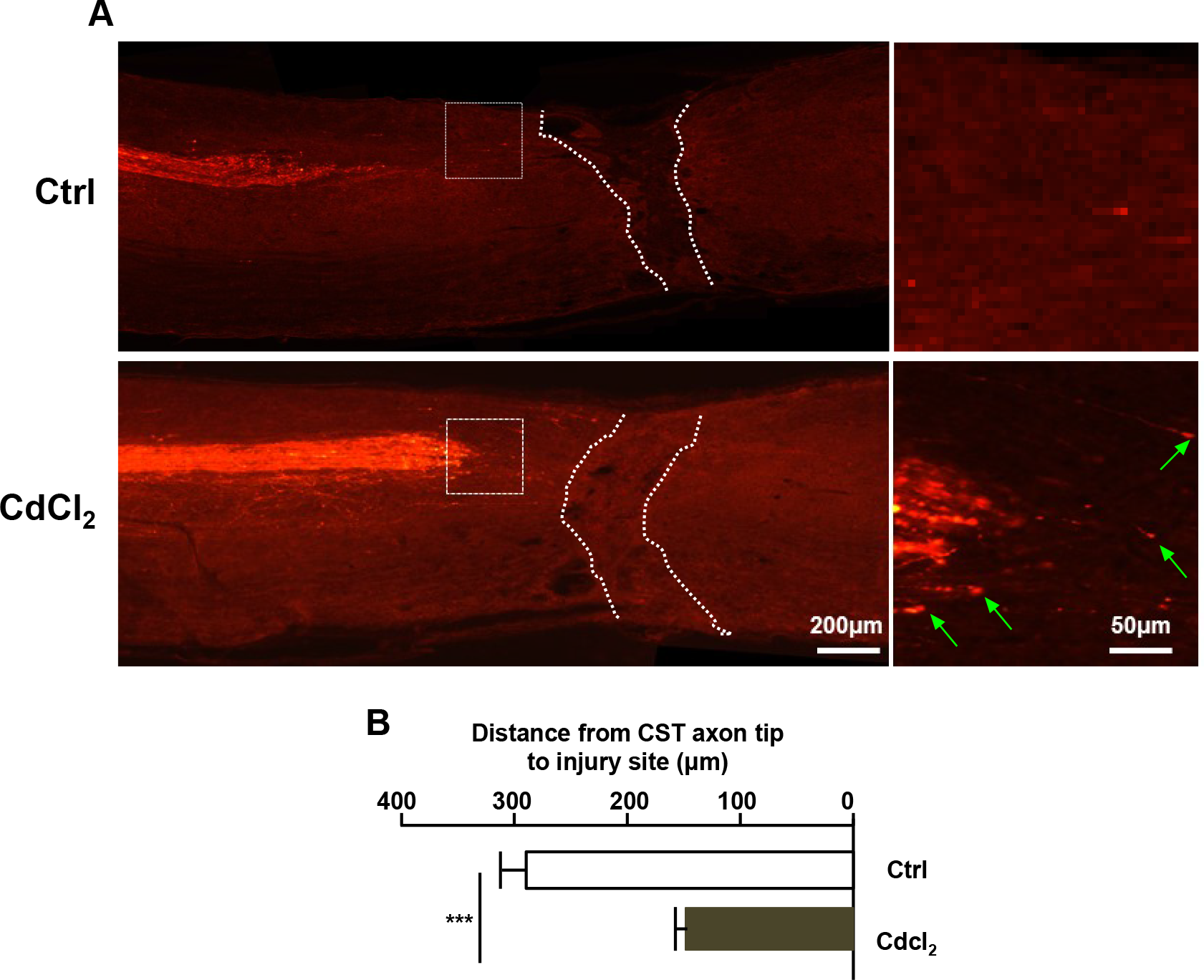
Activation of CaMKII with CdCl2 prevents injury-induced axon retraction of the cortical spinal tract. (A) Representative images of sagittal sections of the spinal cord from intra-peritoneal injection of CdCl2 or vehicle every day for 4 weeks. White dot line: lesion site; Green arrow: retraction bulbs. Scale bar: 200 μm. (B) Measurement of distances of the nearest axon tip from the injury site showing that CdCl_2_ treatment significantly prevented injury-induced axon retraction.

### 7. CaMKII activity inhibition prevents axotomy-induced transcription independent of axon growth

Previously, we developed the culture-and-replating method to investigate whether regenerative axon growth is gene transcription-dependent or local cytoskeleton assembly-dependent [18]. Using this model system, we further investigated whether the regulatory effect of CaMKII on axon growth is gene transcription-dependent or growth cone cytoskeleton assembly-dependent. Adult DRG neurons were dissociated and cultured for 3 days, following which these three-day-cultured neurons were suspended and replated to initiate axon growth anew. Interestingly, we found that neurons treated with KN93 during the initial three-day-culture period had no effect on newly growing axons after replating (Fig. 7A, B). However, axon growth from neurons that were treated with KN93 after replating was significantly blocked (Fig. 7A, B). This result indicates that KN93 mainly inhibits local cytoskeleton assembly-dependent axon growth. On the other hand, ours and others’ previous studies have shown that peripheral axotomy switches the sensory neurons into a regeneration mode, and this peripheral axotomy-induced axon growth is mainly dependent on growth cone cytoskeleton assembly [18]. Thus, adult mice were first subjected to sciatic nerve transection, and seven days after the nerve injury, the L4/L5 DRGs were dissected and cultured for 24 h in vitro with KN93 treatment. In line with the results of the culture-and-replating method, we found the KN93 also blocks peripheral axotomy-induced cytoskeleton assembly-dependent axon growth of DRG neurons (Fig. 7C, D). Together, our data showed that CaMKII mainly regulates local cytoskeleton assembly at the growth cone during axon regeneration.

**Figure 7:**
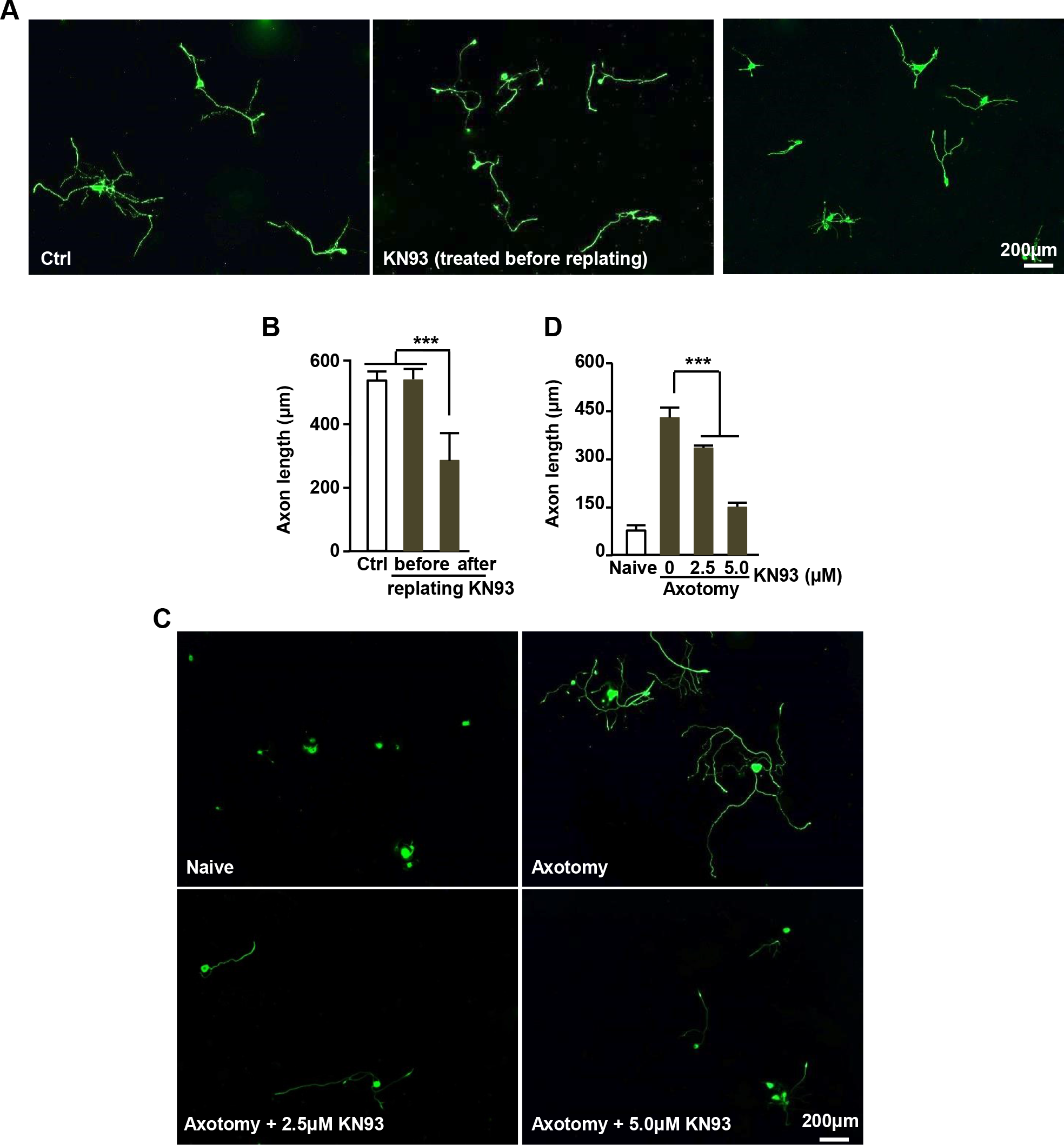
Inhibition of CaMKII activity prevents axotomy-induced axon growth in DRG neurons. (A) Representative images showing only administration of 2.5 μM KN93 after replating blocks axon growth in adult DRG neurons. In contrast, administration of 2.5 μM KN93 before replating did not affect axon growth in adult DRG neurons. Scale bar: 200 μm. (B) Quantification of the average axon length in (A) from three independent experiments. (C) Representative images showing inhibition of CaMKII activity with the pharmacological inhibitor, KN93 (2.5 μM or 5.0 μM), blocks axotomy-induced axon growth of adult DRG neurons. Scale bar: 200 μm. (D) Quantification of the average axon length in (C) from three independent experiments.

### 8. CaMKII regulates the F-actin length in neuronal growth cone

The status of F-actin is known to be regulated by CaMKII in dendritic spines [24]. In addition, our results suggest that CaMKII mainly regulates local cytoskeleton assembly at the growth cone during axon regeneration. Accordingly, we next examined the relationship between the growth cone F-actin status and CaMKII activity. First, our immunoflorescence staining showed that phosphor-CaMKII is located at the frontier tip of the tubulin (Fig. 8A), and co-localized with F-actin (Fig. 8A). Furthermore, we found that inhibition of CaMKII activity with KN93 significantly reduced the F-actin length (Fig. 8B, C). In contrast, activation of CaMKII with cadmium chloride increased F-actin length (Fig. 8 B, C). Thus, it is probable that CaMKII regulates mammalian axon growth by affecting F-actin length in the growth cone.

**Figure 8:**
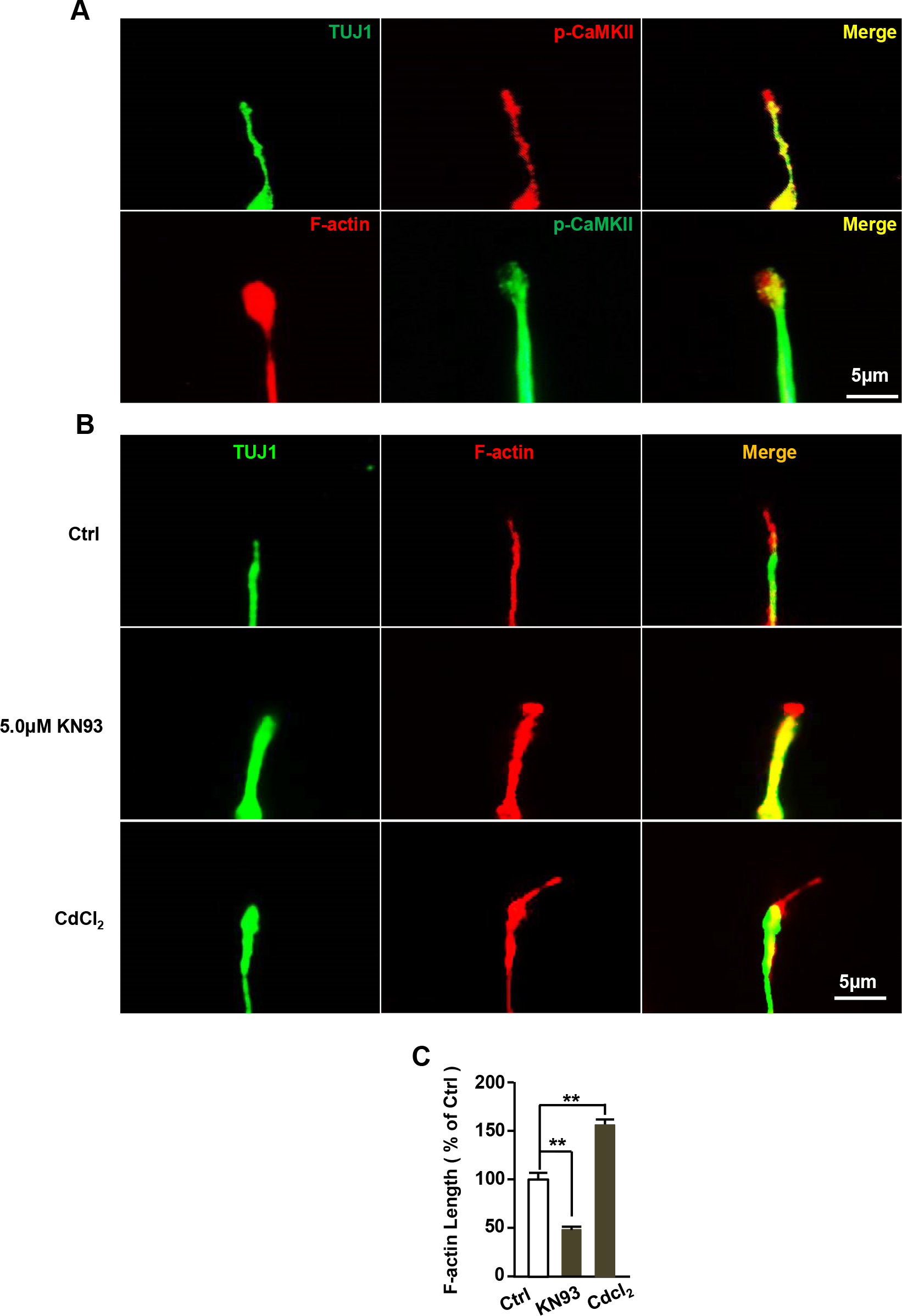
CaMKII affects the F-actin length in growth cone. (A) Immunofluorescence staining showing that phosphor-CaMKII is located at the frontier tip of the tubulin, and co-localizes with F-actin in the DRG neuronal growth cones. Scale bar: 5 μm. (B) Representative images of F-actin staining after treatments of KN93 or CdCl_2_. Scale bar: 5 μm. (C) Quantification of F-actin length showing that inhibition of CaMKII activity with KN93 significantly reduces the F-actin length; in contrast, activations of CaMKII with CdCl2 increase the F-actin length.

## Discussion

The global rates of nerve injury on account of trauma are exceedingly high, and loss of axonal continuity is a common problem associated with functional deficit. Thus, axon regeneration is a key determinant of functional recovery in the injured nervous system. Thus far, several molecules and signaling pathways have been identified that play a potential role in axon regeneration. However, the process of axon regeneration in the mammalian nervous system is very complex. The regenerative capacity of injured neurons is governed by both extrinsic and intrinsic factors. Therefore, axon regeneration in the mature mammalian nervous system is often poor, especially in the CNS neurons.

Herein, we found that peripheral axonal injury increases the phosphorylation level of CaMKII in DRG neurons of adult mice. CaMKII is one of the most prominent protein kinases, and calcium/calmodulin can bind to CaMKII and activate it through autonomous phosphorylation [25]. A previous study has shown that the influx of extracellular Ca^2+^ is one of the first phenomenon that occurs following axonal injury [17]. Thus, it is possible that a peripheral axotomy-induced increase in the concentration of intracellular calcium ions activates CaMKII in DRG neurons. In addition, one study reported that inhibition of calcium influx will reduce the regenerative axon growth in the sensory neurons of *Caenorhabditis elegans* [14]. Similarly, the results of our pharmacological or genetic silencing study showed that inhibition of CaMKII activity impairs mammalian axon regeneration both in vitro and in vivo. More importantly, we also found that activation of CaMKII activity is sufficient to promote embryonic CNS and adult PNS neuronal axon growth in vitro. Additionally, the axons of injured CNS neurons usually form retraction bulbs with a disorganized network of growth cone cytoskeleton elements such as microtubules [20]. Previous studies have shown that stabilization of microtubules with taxol promotes axon regeneration after optic nerve and spinal cord injury [8, 9]. Our result also indicates that activation of CaMKII with CdCl_2_ prevents injury-induced axon retraction after spinal cord injury. Thus, our results showed that CaMKII is an important regulator of axon regeneration, and its activity is required for mammalian axon regeneration, both in vitro and in vivo.

It is well known that axon elongation is achieved by cytoskeletal elements and assembly of membrane components at the nerve growth cone. Another study showed that CaMKII was co-located and interacted with F-actin in the growth cone [26]. CaMKIIβ also contains a binding region for F-actin [27], and regulates its stability [28, 29]. Thus, inhibition of CaMKII causes disorganization of the F-actin structure [26]. On the other hand, CaMKII can phosphorylate many microtubule-associated proteins [30]. For example, CaMKII can inhibit Cofilin functions through LIMK signaling [24], and Cofilin can regulate neuronal actin cytoskeleton by actin depolymerization [31]. Thus, it is possible that CaMKII regulates mammalian axon regeneration by affecting the growth cone cytoskeleton components such as F-actin distribution. First, we found that KN93 mainly inhibits the local cytoskeleton assembly-dependent axon growth, such as peripheral axotomy-induced axon growth. Furthermore, our results showed that F-actin and CaMKII co-localize at the tips of microtubules in the growth cone. Inhibition of CaMKII with KN93 significantly decreases the F-actin length in the growth cone. Conversely, activation of CaMKII with CdCl2 markedly increases the F-actin length in the growth cone. Thus, our data suggested that CaMKII regulates mammalian axon regeneration by affecting F-actin distribution in the growth cone.

In conclusion, our results indicated that peripheral axotomy increases the phosphorylation level of CaMKII in DRG sensory neurons. Moreover, inhibition of CaMKII activity impairs PNS and CNS axon growth in vitro, and activation of CaMKII promotes PNS and CNS axon growth in vitro. Most importantly, our data also indicate that CaMKII activity involves peripheral sensory axon regeneration and cortical tract regeneration in vivo. Furthermore, we also found that CaMKII mainly regulates F-actin length in the growth cone during axon regeneration. Thus, taken together, these findings provide clear evidence that CaMKII is a critical modulator of mammalian axon regeneration.

## Materials and Methods

### Animals and surgical procedure

Thirty 8–10-week-old adult and six pregnant ICR mice were used. All animal experimental procedures were performed according to the animal protocol approved by the Institutional Animal Care and Use Committee of Soochow University. For surgical procedures, all mice were anesthetized with a mixture of ketamine (100 mg/kg) and xylazine (10 mg/kg) via intra-peritoneal (IP) injection. Eye ointment containing atropine sulfate was applied to protect the cornea during surgery, and animals received antibiotics for 24 h as post-operative analgesia.

### Reagents and antibodies

KN93 and Blebbistatin were obtained from Selleck Chemicals. CdCl_2_ was from Sigma-Aldrich. The anti-βIII tubulin (TUJ1) antibody was from Sigma-Aldrich. Primary antibody against β-actin and phospho-CaMKII were from Cell Signaling Technology. Alexa fluor^®^ plus 488 or 568 secondary antibodies were purchased from Life Technologies. Dextran biotin (BDA) was purchased from Life Technologies. Cy3-Streptavidin was from Jackson Laboratories. Actin-stain 555 Fluorescent Phalloidin was from Cytoskeleton. The siRNA against CaMKIIγ was from GenePharma (Shanghai, China) which targeted the following sequences: AACGTGGTACATAATGCTACA, CACAGTCACTCCTGAAGCTAA and ATCATTAAGATCACAGAACAA.

### Cell culture and in vitro electroporation

Cell culture were performed as described in our previous study [32]. DRGs were isolated from 8–10-week-old adult mice and incubated with collagenase (1 mg/mL) at 37℃for 90 min, followed by 0.25% trypsin for 20 min. The DRG neurons were cultured in MEM medium containing 5% fetal bovine serum, 100 mg/mL penicillin and streptomycin. The embryonic cerebral cortical neurons isolated from embryos at E14.5, and hippocampal neurons were isolated from embryos at E18. Then, the embryonic cells were treated with 0.25% trypsin for 5 min and cultured in neurobasal medium supplemented with penicillin/streptomycin, GlutaMAX, and B27 supplements. Either adult DRG or embryonic dissociated cells were cultured on coverslips coated with poly-D-lysine (100 μg/mL) and laminin (10 μg/mL) mixture.

The siRNA were transfected via electroporation according to the manufacturer’s protocol. The dissociated neurons were centrifuged to discard the supernatant, and re-suspended in 100 μL mixture of siRNA or/and EGFP and electroporation buffer. Then, it was transferred to a 2.0-mm electroporation cuvette and electroporated with an Amaxa™ Nucleofector^TM^ device. After electroporation, the cells were immediately mixed with the 500-mL pre-warmed culture medium and plated into a 24-well plate that was coated with poly-D-lysine (100 μg/mL) and laminin (10 μg/mL). After 4 hour, when the neurons had adhered to the coverslips, the medium was replaced to remove residual electroporation buffer.

### Real-time quantitative polymerase chain reaction (RT-PCR)

For RT-PCR, total RNAs were isolated with Trizol reagent (Invitrogen, Carlsbad, CA, USA) and reverse transcribed using Maxima H Minus Reverse Transcriptase (Thermo Scientific, Waltham, MA, USA). Quantitative PCR was performed using SYBR-Green Real-Time PCR Master Mix (Toyobo Co.; Osaka, Japan), and standard curves (cycle threshold values versus template concentration) were prepared for each target gene and the endogenous reference (18S) in each sample. The following primers were used:

CaMKIIγ forward:5′-TACAGTGAAGCTGATGCCAG-3′;
CaMKIIγ reverse: 5′-TTGACACCGCCATCTGACTT-3′

### Western blotting

Tissues or cultured neurons were lysed with RIPA buffer for 30 min at 4°C. The protein concentration was measured by using the BCA Kit, and equal amount of extracted proteins were separated with 10% sodium dodecyl sulfate-polyacrylamide gel electrophoresis (SDS-PAGE). Next, the protein was transferred to a polyvinylidene difluoride (PVDF) membrane (Immobilon-P; Millipore). The membrane was blocked with 5% skimmed milk and incubated with primary antibodies overnight at 4°C, followed by incubation with the appropriate HRP-conjugated secondary antibody at room temperature for 1–2 hour. Afterwards, ECL Prime Western Blotting Detection Reagent (GE Healthcare; Chicago, IL, USA) was used to develop the membrane. ImageJ software was used to quantify the density of protein bands from three independent experiments.

### Immunofluorescence staining

For cultured neurons, the cells were fixed with 4% paraformaldehyde for 20 min and then blocked with 2% BSA and 0.1% Triton X-100 for 1 h, followed by incubating with primary antibody (anti-βIII-tubulin, 1:1000; phospho-CaMKII, 1:500) for 2 h, and secondary antibody for another 1 hour. To visualize the F-actin in growth cone, the DRG neurons were incubated with Actin-stain 555 fluorescent Phalloidin for 30 min. Then, the samples were again washed thrice in PBS and mounted with Mowiol^®^.

For immunohistostaining, the mice were anesthetized and transcardically perfused with 4% paraformaldehyde under deep anesthesia. The L3-L4 DRG tissue was dissected out and further fixed with 4% paraformaldehyde overnight at 4°C. The tissues were cryo-sectioned at 12-μm thickness. Then, the sections were blocked in PBS containing 10% FBS and 0.3% Triton X-100 for 1 h and incubated over-night with anti-βIII-tubulin (TUJ1, 1:1000) antibody and phospho-CaMKII (1:500) antibody. Next day, the sections were washed thrice in PBS, and then incubated with fluorescent-labeled secondary antibody for 1 h. The samples were again washed thrice in PBS and mounted with Mowiol^®^.

### In vivo DRG electroporation

In vivo electroporation was performed as described in our previous study [32]. After mice were anesthetized and the left side of the L3-L4 DRGs was exposed, a 1-μl solution containing siRNA and/or GFP plasmid was carefully micro-injected into each DRG (pressure: 30 psi, duration: 8 ms) using a capillary pipette powered by Picospritzer Ⅲ (Parker Inc., Cleveland, OH, USA). Electroporation was performed immediately after microinjection using tweezer-like electrodes (Ø1.0 mm) and ECM830 electric transducers BTX (35 V, 15-ms pulse, and 950-ms interval). The skin was subsequently sutured. Two days later, a sciatic nerve crush injury was created on the isolateral side using fine forceps, and the injury site was marked with a 11-0 nylon epineural suture. After another three days, the mice were perfused with 4% paraformaldehyde, and the entire sciatic nerve was dissected out. Using AxioVision 4.7 software (Carl Zeiss MicroImaging, Inc.), the lengths of all EGFP-positive axons were measured manually from the crush site to the distal axon tips in whole-mounted tissue.

### Measure of axon length and F-actin length

All images were captured with CCD camera controlled by AxioVision 4.7 software (Carl Zeiss MicroImaging, Inc.), and axon length was measured using the “measure/Curve/Outline” application. To quantify the axon length, neurons with axons more than twice the diameter of the cell body were counted and the longest axons of each neuron were tracked manually using the “measure/Curve/Outline” application. More than 100 neuronal axons were measured in three independent experiments. For quantification of F-actin length in the growth cone, we measured the distance from the fluorescent phalloidin-labeled F-actin tips to TUJ1 labeled microtubules ends with the AxioVision 4.7 software.

### Corticospinal tract tracing and axon retraction measuring

To trace the cortical spinal tract, we injected BDA into the sensorimotor cortex as indicated indicated previously [33]. In brief, 14 days post injury, a total of 1.6 μl 10% BDA in PBS was injected into the sensorimotor cortex of four sites (1.0 mm lateral; 0.5 mm deep into the cortex; and 1.0, 0.5, −0.5, and −1.0 posterior from the bregma) with a 5-μl Hamilton micro syringe. Fourteen days after injection, mice were perfused with 4% PFA. Spinal cords were isolated and further fixed in PFA for 6 h at 4°C. After dehydration in 30% sucrose, tissues were sectioned either sagittal or cross sectioned to give 25-μm thick sections. Sections of spinal cords were stained with Cy3-Streptavidin (Jackson, 1:500) for 3 h. After washing with PBS thrice, sections were mounted with mounting solution to obtain photos using fluorescence microscopy. To measure axon retraction, distances from the axon tips to the center of the lesion site were measured. The averages of all axons of five sections the spinal cord were calculated and at least five mice were used for each group.

### Statistics

Data are expressed as mean ± standard error of the mean (SEM). “n” represents the number of independent experiments. The significant differences between groups were determined by single factor with *t*-test. Statistical significance was set to p<0.05 (*: p<0.05; **: p<0.01; ***: p<0.001).

## Author Contributions

F X, RJ X, JH X, WH W, JJ M, F W, YX M, SB Q, JC Z, HN Z, XZ Q, JQ C, B L, CM L, HL Y, B M, and Saijilafu designed the experiment. F X, JH X and WH W performed the experiments and analyzed the data. F X, RJ X and Saijilafu co-wrote the paper with all authors’ input.

## Acknowledgements

This work was supported by a grant from the the National Key Research and Development Program (No. 2016YFC1100203), National Natural Science Foundation of China (No. 81571189 and No. 81772353 to Saijilafu), A Priority Academic Program Development of Jiangsu Higher Education Institutions, and An Innovation and Entrepreneurship Program of Jiangsu Province. Science & Technology Development Program of Suzhou City (to B M).

